# The Golgi matrix protein giantin is required for normal cilia function in zebrafish

**DOI:** 10.1101/117507

**Authors:** Dylan J.M. Bergen, Nicola L. Stevenson, Roderick E.H. Skinner, David J. Stephens, Chrissy L. Hammond

**Affiliations:** Cell Biology Laboratories, School of Biochemistry, University of Bristol, Biomedical Sciences Building, University Walk, Bristol, UK, BS8 1TD; School of Physiology, Pharmacology and Neuroscience, University of Bristol, Biomedical Sciences Building, University Walk, Bristol, UK, BS8 1TD

**Keywords:** Golgi, extracellular matrix, glycosylation, cilia, zebrafish

## Abstract

The Golgi is essential for glycosylation of newly synthesised proteins including almost all cell-surface and extracellular matrix proteoglycans. Giantin, encoded by the *golgb1* gene, is a member of the golgin family of proteins that reside within the Golgi stack but its function remains elusive. Loss-of-function of giantin in rats causes osteochondrodysplasia; knockout mice show milder defects, notably a cleft palate. In vitro, giantin has been implicated in Golgi organization, biosynthetic trafficking, and ciliogenesis. Here we show that loss-of-function of giantin in zebrafish, using either morpholino or knockout techniques, causes defects in cilia function. Giantin morphants have fewer cilia in the neural tube and those remaining are longer. Mutants have the same number of cilia in the neural tube but these cilia are also elongated. Scanning electron microscopy shows that loss of giantin results in an accumulation of material at the ciliary tip, consistent with a loss-of-function of retrograde intraflagellar transport. Mutants show milder defects than morphants consistent with adaptation to loss of giantin.

**Summary statement:** Loss of giantin following either morpholino injection or genome editing in zebrafish results in defects in ciliogenesis.

## Introduction

The Golgi apparatus is the main hub of the secretory pathway, responsible for modifying proteins derived from the endoplasmic reticulum (ER) prior to transportation to the plasma membrane. It is estimated that one-third of the proteome traverses the ER-to-Golgi transport pathway (Zanetti et al., 2011). This includes delivery of key developmental signalling receptors to the plasma membrane and assembly of the extracellular matrix (ECM), which present a major secretory load during development of the early embryo (Zhong, 2011). The organisation of Golgi membranes into flattened disc shapes is orchestrated by eleven members of the Golgi matrix protein family (golgins) that decorate the Golgi surface to form a mesh. Many of these proteins mediate the docking and tethering of coat protein (COP) coated transport vesicles, COPI and COPII, at the Golgi cisternae, whilst others are involved in inter-cisternal tethering (Gillingham and Munro, 2016). These functions are carried out by coiled-coiled domains common to all golgins, which autonomously fold into long rod-like structures. Golgins are sufficient to direct the tethering of incoming vesicles to the Golgi with some overlap seen in their function (Wong and Munro, 2014).

The largest member of the golgin family is giantin (encoded by the *golgb1* gene), which has a single C-terminal transmembrane domain (TMD) and 37 predicted coiled-coiled domains. As such, its α- helical coils have a predicted reach of up to 450 nm into the cytoplasm from the *cis-and-medial* Golgi membranes (Munro, 2011). The exact function of giantin is still unclear. There is biochemical evidence of a role for giantin in tethering COPI vesicles (Yoshimura et al., 2004), however artificial relocation of giantin to mitochondria does not reveal a clear role in tethering incoming vesicles (Wong and Munro, 2014). This suggests that either it does not act as a tether or that its function is redundant with that of other golgins. Giantin is present in all vertebrate genomes but is not found in *C. elegans* or *Drosophila.* In those organisms that express it, giantin is widely expressed and is found in almost all tissues.

Primary cilia are microtubule-based apical cell projections, built as extensions from the mother centriole in almost all non-cycling cells. Ciliary cargo, varying from cilia specific proteins to signalling pathway components, is trafficked along the microtubule-based axoneme by intraflagellar transport (IFT). Anterograde movement from the base to the tip is driven by IFT-B/kinesin-2, whereas retrograde transport is IFT-A/dynein-2 driven (Garcia-Gonzalo and Reiter, 2012). This equilibrium between anterograde and retrograde IFT is crucial for cilia to act as signalling platforms for many signalling pathways including some key to development such as the Hedgehog (Hh) signalling cascade. Members of the IFT complex and Hh signalling cascade are mutated in ciliopathies associated with severe skeletal defects including short rib polydactyly and Jeune syndrome (Huber and Cormier-Daire, 2012).

Our previous work revealed an unexpected role for giantin in ciliogenesis (Asante et al., 2013). We showed that acute depletion of giantin in epithelial cells results in fewer but longer cilia (Asante et al., 2013). Giantin is required for the localisation of the dynein-2 intermediate chain WD-repeat protein 34 (WDR34) at the base of the cilium. Since dynein-2 is the retrograde motor for IFT within cilia, we reasoned that loss of giantin might affect transport within the cilium and explain the defects in both ciliogenesis and length control.

Rodent models with loss-of-function mutations in giantin have been described. A spontaneous 10bp insertion in *Rattus norvegicus golgb1* (Rn-Golgb1) was shown to be a functional knockout of giantin. This recessive mutation leads to an osteochondrodysplasia (ocd) phenotype with homozygous mutant pups exhibiting craniofacial abnormalities, oedema, shorter limbs, and defects in collagen-rich matrix deposition, leading to late embryonic lethality (Katayama et al., 2011). More recently, *golgb1* knockout mice were generated using both N-ethyl-N-nitrosourea (ENU) and CRISPR/Cas9 mutagenesis (Lan et al., 2016). These animals show no gross abnormalities in skeletal development, including of craniofacial structures, other than cleft palate and an abnormal deposition of glycosylated bone ECM as recognised by peanut agglutinin (PNA) lectin (Lan et al., 2016).The discrepancies in severity of the phenotypes of these rodent models could be due to species differences or genetic background. Nonetheless, both show extensive defects in ECM secretion and/or assembly.

Here, we describe that acute depletion of giantin in zebrafish leads to defects in cilia number and structure. These data show that acute depletion of giantin *in vivo* recapitulates ciliary defects seen *in vitro.* They also suggest that adaptive changes might underlie the phenotypic variability of giantin knockout animals.

## Results

### Acute knockdown and genetic knockout approach to assess giantin function in the zebrafish

Zebrafish giantin is encoded by 20 exons, with the translational start site (TSS), the human to zebrafish conserved coding sequence for the p115 binding site, and C-terminal TMD at exon 2, 3, and 20 respectively (Figure 1A). The relative exon position of existing mutations identified or generated in rat and mouse models (Katayama et al., 2011; Lan et al., 2016), shows that all mutations affect evolutionary conserved exons, ranging from exon 10 to 15 (Figure 1A, yellow, blue, and brown coloured bars).

**Figure 1:**
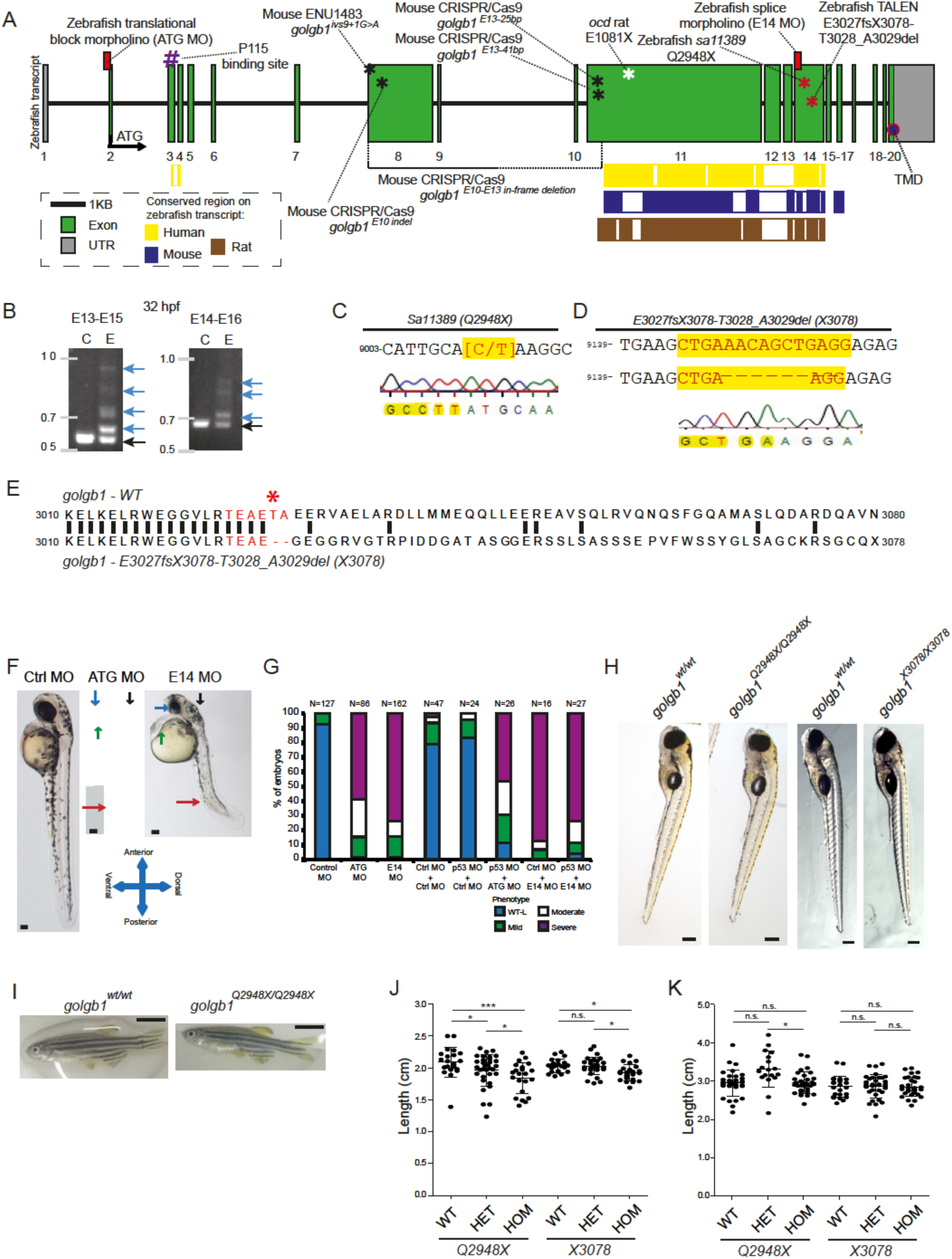
Experimental knockdown and knockout of giantin *in vivo*. (A) Schematic representation of the zebrafish *golgb1* transcript (ENSDART00000131402.2) showing the binding sites of the designed morpholinos (ATG and E14), the relative exonic location of the human annotated P115 binding site, transmembrane domain (TMD), the various mouse mutants (black asterisks) and the *ocd* rat allele (white asterisk) as described in Lan Y *et al.* 2016, and Katayama K *et al.* 2011 respectively. Coloured bars show the interspecies conserved sequence regions as assigned in Ensembl (release 87). (B) Alternative spliced PCR products (blue arrows) from 32 hpf RT-PCR cDNA of two amplicons in E14 MO binding region. (C) *Sa11389* line carrying a point mutation from C to T (yellow highlight, red letters) resulting in a premature stop codon at Q2948 *(golgb1^Q2948X^)* from the EZRC. (D) TALEN site directed mutagenesis resulted in an 8bp deletion (spacer sequence in yellow, red letters), resulting in *golgb1^X3078^* mutant line. (E) Alignment of Golgb1 WT protein sequence (Ensembl release 87) with predicted *golgb1^X3078^* protein sequence showing translated spacer sequence (red) and site of deletion T3028 and A3029 with a subsequent frameshift from E3027 (red asterisk) changing 51 amino acids to a predicted stop codon at position 3078. Part of cDNA exon 14 sequence. (F) Stereomicroscope images of 48 hpf *control, ATG,* and *E14* morphants exhibiting defects in the eye, heart, cranium, and various axis orientations (blue, green, black, red arrow respectively). (G) Percentile quantification of scored phenotypes. (H) Stereomicroscope images of heterozygote incross 5 dpf larvae from both mutant lines and (I) 7 wpf female adults. (J) Dot plot for body length at 7 wpf, and (K) 44-46 wpf for *golgb1^Q2948X^* and 41-43 wpf for *golgb1^X3078^* alleles. Scale: (F) 100 μm, (H) 200 μm, (I) 500 μm. (J, K) 1-way ANOVA with Tukey's Multiple Comparison Test. All experiments of three replicates. Bars show means with standard deviation. * p<0.05, ** p<0.01, and *** p<0.005.

We chose to analyse the role of giantin in zebrafish using a combination of morpholino knockdown and genetic mutants. To knockdown giantin, two antisense morpholinos (MOs) were designed against the zebrafish *golgb1* gene, the first targeting the TSS (ATG MO) located in exon 2 (ATG MO Fig 1A) and a second targeting a highly-conserved splice site located across intron 13 and the start of exon 14 (E14 MO, Figure 1A). We validated the efficacy of the E14 MO using reverse transcription (RT)-PCR on cDNA generated from RNA extracted from controls and E14 morphants. Abnormal splicing following injection with the E14 morpholino was demonstrated by RT-PCR (Figure 1B). To complement our MO knockdown experiments, we also examined the role of giantin using two mutant alleles. The sa11389 mutant, carries a (C>T) point mutation in exon 14 (Figure 1A and 1C) resulting in generation of a premature stop codon at glutamine-2948 (henceforth denoted *golgb1^Q2948X^).* In addition to the ENU allele we generated a new mutant line by TALEN mutagenesis, whereby TAL arms were targeted to a conserved region in exon 14 (Figure 1A). Injection of capped mRNA including these arms gave rise to genetically mosaic fish, and by screening the F1 generation we identified a carrier with an 8bp insertion in exon 14 in the spacer sequence (Figure 1C). This resulted in a frameshift at position 3029, and a premature stop codon at position 3078 (E3027fsX3078-T3028_A3029del, henceforth called *golgb1^X3078^)* (Figure 1D and 1E). This carrier transmitted the identified mutation through the germline allowing a mutant line to be maintained. Both lines are predicted to be loss-of-function mutations as they include premature stop codons that terminate the coding region upstream of the transmembrane spanning region at the C-terminal end of the protein.

Embryos injected with either the ATG or the E14 MO show no obvious morphological defects for the first 24 hours of development, however by 2 days post fertilisation (dpf) larvae display a range of phenotypes including smaller eyes, reduction in total body length, kinks and curls in the tail and frequent oedema of the heart and brain (Figure 1F). These phenotypes are characteristic of ciliopathies in zebrafish. We characterised relative severity based on phenotypical characteristics depicted in giantin morphants into four groups: wildtype-like (WT-L), mild, moderate, and severe. The mild classification was assigned when one or two characteristics were observed in a mildly manner. Identifying three characteristics of which at least one in a severe manner, or four 'mild' phenotypes were classed as moderate. Severe labels were assigned to individuals exhibiting 4 or more phenotypical outcomes, of which at least two severe at 2 dpf. The relative distribution of phenotypes between ATG and E14 MOs were similar at 48 hpf. Moreover, co-injection with p53 MO did not alter the phenotypes observed in giantin MOs (Figure 1G), implying that the smaller embryo size was not caused by p53-induced apoptosis.

To determine whether the phenotypes seen on acute knockdown of giantin using morpholinos would be recapitulated in a knockout we studied the larvae of the two loss-of-function mutant alleles described above. Somewhat surprisingly, these homozygous giantin knockout fish did not show any major morphological defects at 5 dpf (Figure 1H). Interestingly, homozygous mutants carrying either allele are viable (Figure 1H) and can produce viable offspring. During juvenile development, they did show a small but significant difference in growth (a reduction of 11.86% and 5.4% respectively along the anterior to posterior axis) at 50 dpf (Figure 1I, 1J). At 44-46 weeks post fertilization (wpf) for *golgb1^Q294SX^* and 41-43 wpf for *golgb1^X3078^*alleles, the reduction in body length was not evident (Figure 1K) indicative of a delay rather than a defect in growth.

### Knockdown of giantin leads to a ciliopathy-like phenotype, longer and fewer cilia, and alterations in cilia morphology

Our previous *in vitro* work demonstrated a role for giantin in cilia function (Asante et al., 2013). To assess the role that giantin plays in ciliogenesis *in vivo,* we visualised cilia in the ventral neural tube of morphants at 24 hpf by labelling for acetylated tubulin (Figure 2A). Both ATG and E14 injected morphants showed a significant reduction in cilia number (decreased by 22% and 33.5% respectively, Figure 2B) with a significant increase in length of remaining cilia (Figures 2C and 2D showing a 22.8% and 34.5% increase in ciliary length for *ATG* and *E14* morphants respectively). Moreover, imaging of the ciliary membrane marker Arl13b tagged to GFP (Arl13b-GFP), showed longer cilia in transgenic E14 morphant trunks at 3 dpf (Figure 2E-G).

**Figure 2:**
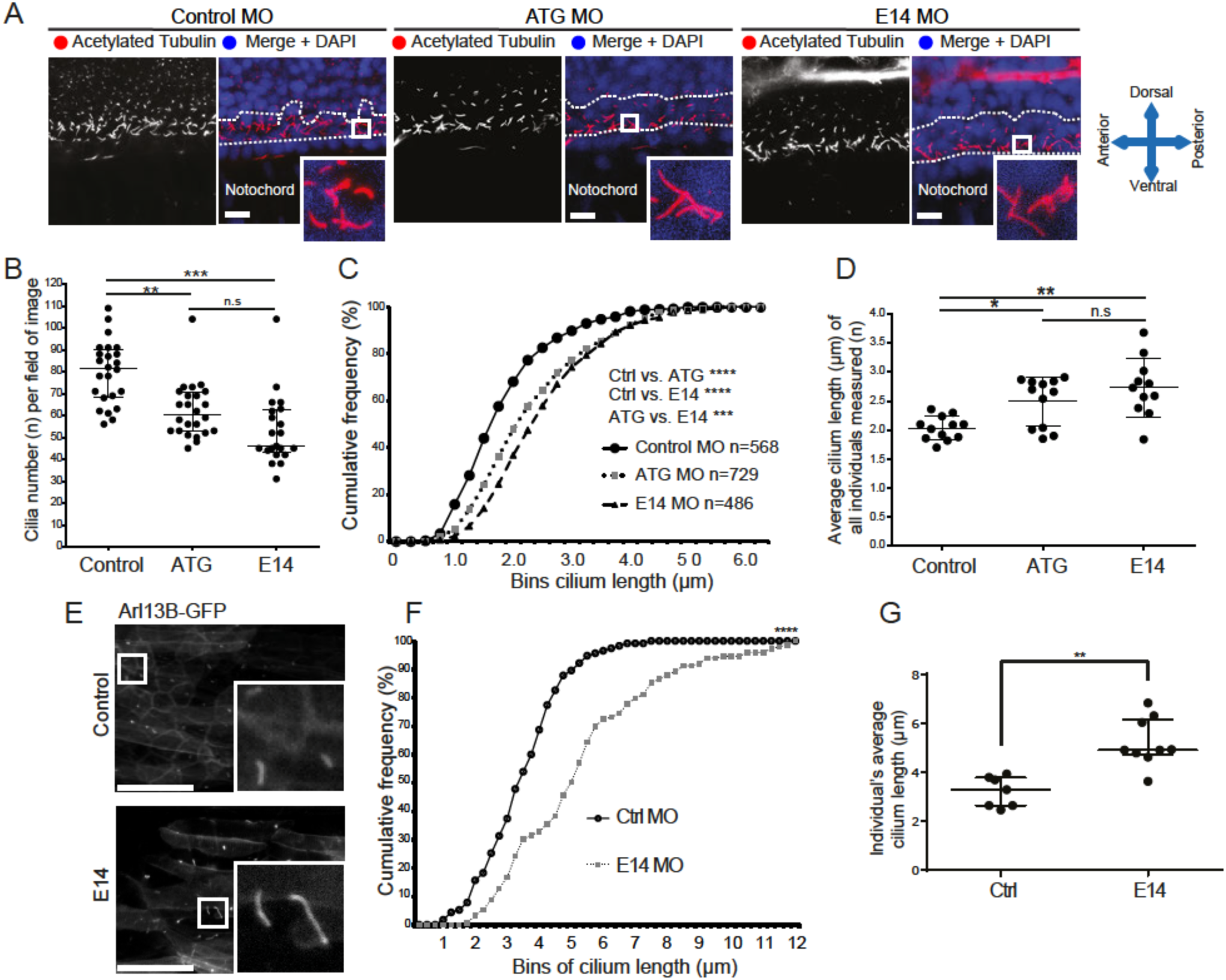
Giantin knockdown leads to changed cilia length and morphology with a ciliopathy-like phenotype in morphant zebrafish. (A) Lateral view confocal images of 24 hpf neural tube ependymal cilia (red in merge) were quantified in a two nuclei wide area (dotted line). Nuclei (DAPI-labelled) are shown in blue. Bar = 10 μm. (B) Quantification of cilia number per field of image within the two nuclei window (panel A, two images per individual). Bar = 50 μm. (C) Cumulative frequency plot of all cilia measured. (D) Average cilium length per individual measured. (E) Representative confocal images of *in vivo* imaging control and E14 morphants showing Arl13b-GFP positive cilia in the myotome (lateral view) Scale: 10μm. (F) Arl13b-GFP ciliary membrane length displayed in a cumulative frequency plot. Data from 2 independent experiments. (G) Average cilium length per individual imaged, asterisk shows significance assessed by t-test. On scatter plots (B, D), bars represent mean with standard deviation and statistical testing was done by one-way ANOVA with a Dunn's multiple comparison test. * p<0.05, ** p<0.01, and *** p<0.005.

We then examined cilia in giantin KOs (Figure 3B, compared to controls, Figure 3A). In contrast to morphants, no significant difference was seen in cilia number in the neural tube of giantin knockouts compared to controls (Figure 3C). Quantifying cilia length in both *golgb1^Q2948x/Q2948x^*and *golgb1^X3078/X3078^* mutants at 1 dpf showed a reproducible increase in ciliary length in the ventral neural tube (15.9% and 19.8% respectively) compared to WT (Figures 3D, showing average cilia length per individual and 3E showing cumulative frequency plots of pooled data from multiple fish).

**Figure 3:**
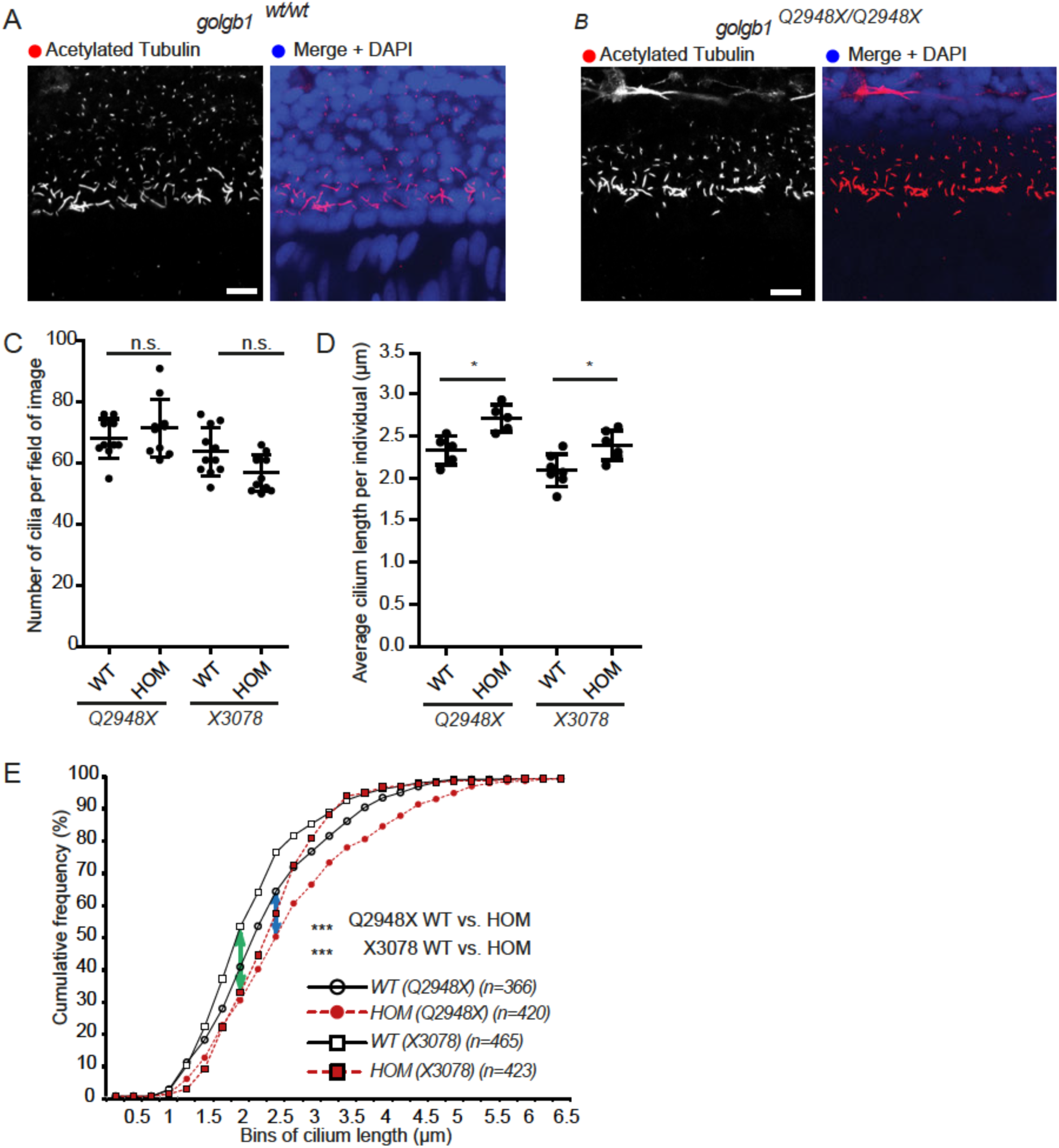
Knockout of giantin results in defects in cilia length. (A, B) Lateral view confocal images of 24 hpf cilia in ventral neural tube of wildtype (A) and homozygous mutant (B). Bar = 10 μm. (C) Cilia number per field of image (two per individual) in a two nuclei wide area adjacent to the notochord. (D) Average ciliary length per individual imaged (confocal). Data shown from at least three independent injection rounds. Bars show mean with standard deviation. One-way ANOVA with a Dunn's multiple comparison test was performed. * p<0.05 (E) Cumulative frequency plot showing shift (blue arrow *Q2948X* and green arrow *X3078)* in percentile abundance of binned (0.25 μm bins) ciliary length between WT and homozygous 24 hpf embryos.

Since we saw an increase in cilia length in both morphants and mutants *in vivo* upon loss of giantin we wanted to assess the appearance of the cilia. At 3 dpf, the olfactory placode is located at the anterior side of the head and projects cilia into the lumen for sensory purposes. Imaging of olfactory pit cilia using scanning electron microscopy (SEM) showed that *control* morphants had a high density of olfactory cell cilia with a uniform appearance along their length, including at the tip. In contrast, both *ATG* and *E14* morphants displayed a lower density of these cilia and an obvious bulbous appearance to the ciliary tip (Figure 4A). This phenotype could be reproduced *in vitro,* using the porcine kidney LLC-PK1 cell line transfected with short interfering RNAs (siRNA) targeting giantin led to bulbous ciliary tips compared to the smooth cilia in mock knockdown cells (Figure 4B).

**Figure 4:**
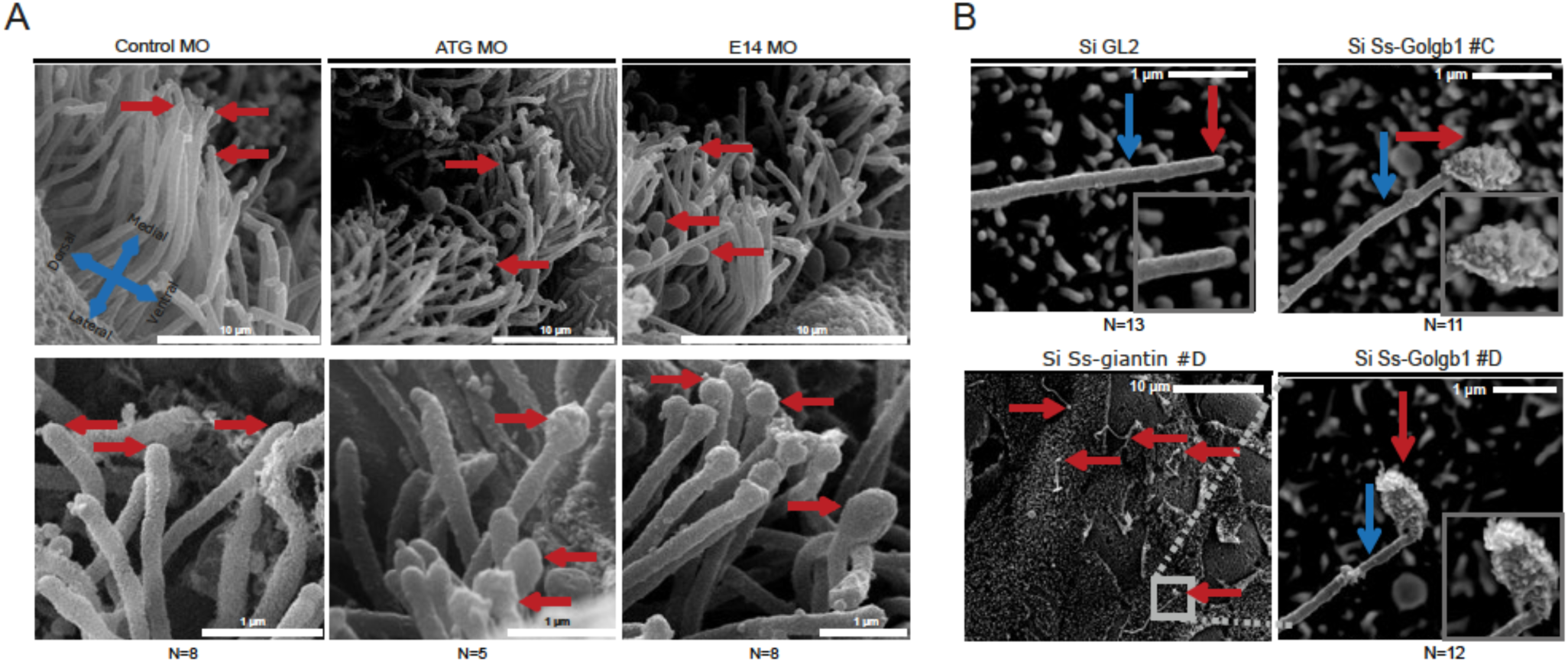
Scanning electron microscopy reveals structural defects in cilia following knockdown of giantin. (A) SEM images olfactory pit (3 dpf) indicating ciliary tips (arrow). (B) SEM images of LLC-PK1 cells exhibit bulbous ciliary tips (red arrow) with visible ciliary extension (blue arrow) after *Ss-Golgb1* knockdown.

Consistent with depletion of giantin causing defects in cilia function, MO knockdown of giantin using either ATG or E14 splice MOs caused heart laterality defects. The heart appeared morphologically normal but was situated either centrally or to the right rather than in its usual left position in a significant percentage of larvae at 32 hpf (Figure 5A). We then sought to rescue the effect of either the ATG or E14 giantin MO by co-injection with mRNA encoding FLAG-tagged giantin from *Rattus norvegicus* (Rn-Golgb1). Rat giantin was seen to localise to Golgi-like structures by immunofluorescence (Figure 5B). Randomisation of the heart position could be partially rescued (ARG: 17% and E14: 43%) by co-injecting the MOs with mRNA encoding Rn-Golgb1 (Figure 5B and 5C). This ability to rescue the phenotype using mRNA from a mammalian species demonstrates that the function of giantin is conserved between rat and zebrafish.

**Figure 5:**
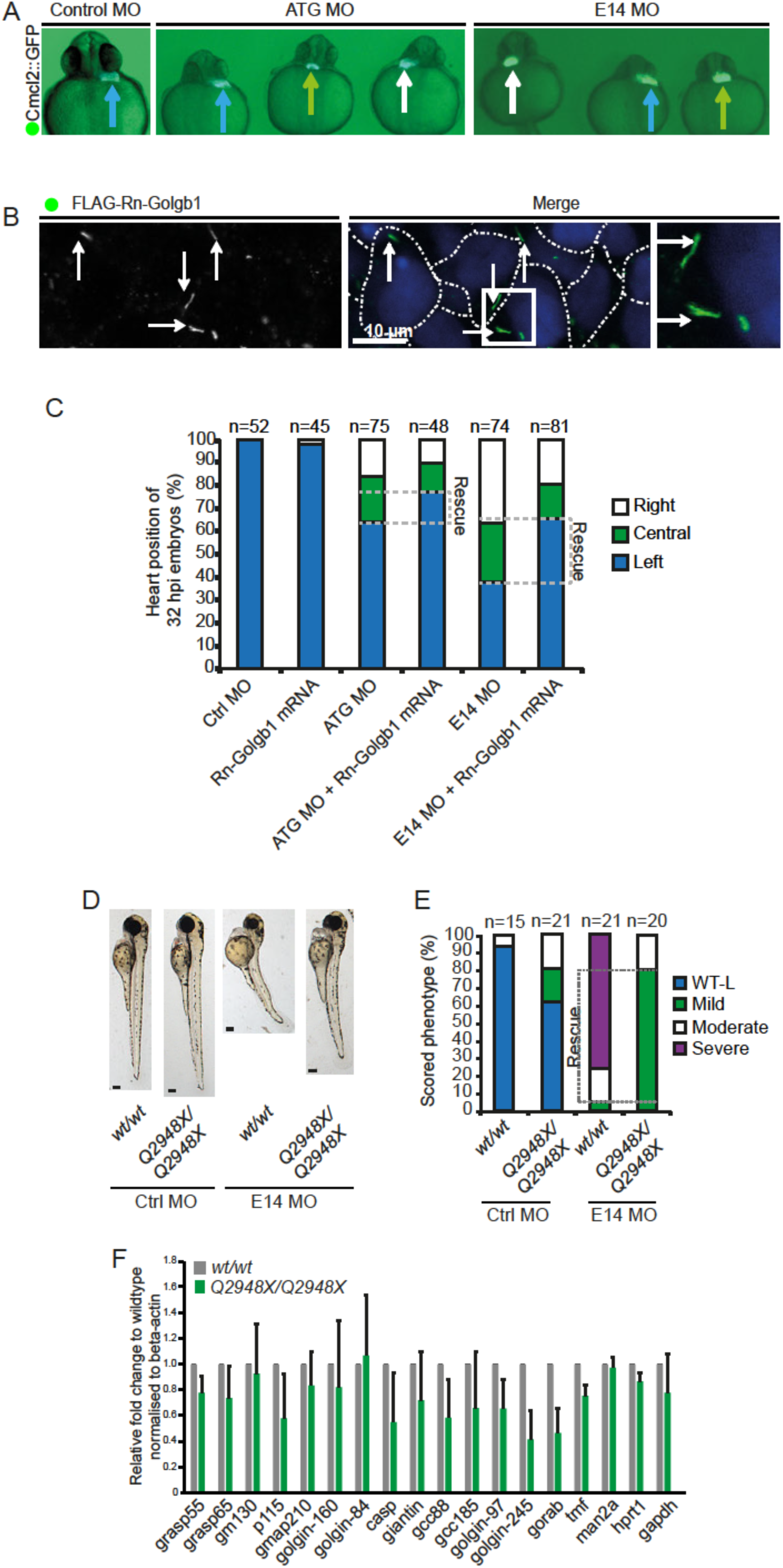
Left-right asymmetry and phenotypic rescue of giantin MO. (A) Cardiomyocyte Cmcl2::GFP transgenic reporter line outlining the heart position (arrow, blue: left; green: midline; white: right) at 32 hpf. (B) Representative confocal image of 48 hpf with ATG morpholino and Rn-Golgb1 mRNA showing Golgi localisation of Rn-Golgb1 in dermal cells stained for anti-flag (white arrow). White dotted line is indicating cell boundaries. N=3 individuals. (C) Quantification of heart position (32 hpf), additionally showing Rn-Golgb1 mRNA co-injection rescue. (D) Stereomicroscope images of 3 dpf control and E14 morpholinos injected in *WT* and *goigb1^Q2948X/Q2948X^* individuals with (E) quantification of phenotypes scored. (F) Quantitative RT-PCR of golgins showing relative mRNA expression levels on 5 dpf larvae. Normalised to beta-actin, mean and standard deviation. All data from three independent experiments.

The availability of a mutant null for giantin enabled us to test the specificity of the E14 MO. We injected the E14 splice morpholino into the Q2984X mutant line. As described in Figure 1, when the MO was injected into WT embryos 73% of larvae exhibited the ‘severe’ ciliopathy-like phenotype. In contrast, mutant embryos injected with the morpholino did not display the most severe phenotypes, instead a majority (77.5%) of the mutants displayed a mild phenotype (Figure 5D, quantified in 5E). Our interpretation here is that the severe phenotypes (as described in Figure 1) are due to depletion of giantin following MO injection and not due to off-target effects.

While knockdown and knockout fish show similar phenotypes, there are clear differences in terms of severity. These differences between acute and chronic loss of giantin are perhaps most easily explained by long-term compensation in the mutant fish via changes to expression of other genes. Since golgins have been shown to have overlapping functions (Wong and Munro, 2014), indicating functional redundancy, we performed a qPCR of golgins (GRASP55, GRASP65, GM130, GMAP-210, golgin-160, golgin-84, CASP, GCC88, GCC185, golgin-97, golgin-245, GORAB, and TMF) and p115, a well-defined giantin interacting protein, to determine whether any were upregulated at the transcriptional level in response to loss of giantin function. No significant difference in golgin expression was found when determining the mRNA expression levels of 15 amplicons from 3 independent total RNA pools of 5 dpf *WT* and *golgb1^Q2948X/Q2948X^* mutant larvae (Figure 5F). This suggests that the compensatory effects seen in the mutants are not mediated by upregulation of other golgins *in vivo.*

A major ciliary pathway linked to developmental patterning is the sonic hedgehog pathway. Smoothened agonist (SAG) treatment could not rescue the phenotypic defects in giantin morphants (Supplemental Figure S1). Consistent with this, smoothened trafficking was also indistinguishable between WT and giantin knockdown hTERT-RPE1 cells (Supplemental Figure S2). Therefore, the phenotypes described here are not likely to result from defective Shh signalling.

## Discussion

Here we show that the largest Golgi matrix protein, giantin, plays a role in ciliogenesis and control of ciliary length *in vivo.* Previous work showed that giantin is required for the ciliary localisation of the WDR34 intermediate chain subunit of the retrograde IFT motor dynein-2 (Asante et al., 2013). Knockdown of giantin in mammalian cells also led to a reduction in cilia number whilst the remaining cilia were longer (Asante et al., 2013). Mutations in human dynein-2 components lead to skeletal ciliopathies (McInerney-Leo et al., 2013; Schmidts et al., 2013a; Schmidts et al., 2015; Schmidts et al., 2013b) where defects in cilia formation and/or function result in pathologies associated with formation of the skeleton (Huber and Cormier-Daire, 2012). These ciliogenesis defects seen in cells are phenocopied here *in vivo* where we show that *golgb1* morpholino knockdown leads to a ciliopathy-like phenotype. Bulbous ciliary tips are seen upon acute depletion of giantin in zebrafish and in cultured pig kidney epithelial cells, consistent with a defect in retrograde IFT. This is also consistent with other work showing loss of the dynein-2 motor heavy chain subunit leads to shorter cilia and bulbous tips in the zebrafish (Krock et al., 2009). In agreement with morphant data, we found that cilia in the neural tube were significantly longer in homozygous mutant fish. This phenotype was not fully penetrant as cilia number was normal, potentially through tissue specific compensation upon chronic loss of giantin.

Some phenotypic discrepancies between giantin morphants and homozygous mutants were observed, suggesting a compensation mechanism following gene knockout. Overall, the effect of giantin knockout is less significant than depletion by morpholino injection. It has been estimated that 90-95% of genomic zebrafish knockouts do not resemble the acute morpholino knockdown phenotypes (Kok et al., 2015). This discrepancy between knockdown and genetic knockout phenotypes has also been shown in other model systems including mouse (Daude et al., 2012; De Souza et al., 2006) and *Arabidopsis* (Gao et al., 2015). Recently, Rossi *et al.* described that 'long term' ECM-dependent compensatory mechanisms are active to compensate for chronic loss of vascular ECM regulator *egfl7.* RNA sequencing screening led to the identification of Emilin2 and Emilin3 ECM components as compensatory factors in genetic *egfl7* knockouts (Rossi et al., 2015). Our work suggests that genetic compensation in the mutant fish is unlikely to have occurred through changes in expression of other golgins, as mRNA transcript levels were equivalent in WT and mutant fish. Owing to a lack of effective reagents, we were not able to determine the effect on protein expression by immunoblot or immunofluorescence. Similarly, we were not able to rescue phenotypes on exogenous stimulation of hedgehog signalling and we could detect no defects in smoothened trafficking to cilia in giantin-depleted cells in culture.

We also note considerable variation in phenotypes between the mouse (Lan et al., 2016), rat (Katayama et al., 2011), and zebrafish *giantin* knockout animal models. Knockout mice have non-syndromic cleft palate but show limited skeletal phenotypes; knockout rats show osteochondrodysplasia. Both rodent models die during late embryonic development. In contrast our fish survive to adulthood. This might reflect differences in developmental pathways, or in the mechanisms of compensation for loss of giantin function. Our previous *in vitro* work implicated giantin in the secretion of ECM (McCaughey et al., 2016) and we are now actively studying this in our knockout model. It is possible that the capacity to adapt the ECM during development varies between both tissues and species and that this could compound the variation in phenotypes observed. This variety of phenotypes may also partially explain why acute knockdown and chronic knockout of *golgb1* show differences, since acute knockdown may not trigger an adequate compensatory response. In summary, our data using *in vivo* models complement our previous *in vitro* work to support a role for giantin in controlling cilia formation and function.

## Material and methods

### Zebrafish husbandry and transgenic lines

The London AB strain was used and maintained according to standard conditions (Westerfield, 2000) and staged accordingly (Kimmel et al., 1995). Transgenic lines: Arl13-GFP fusion protein *(Tg(Act-B2::Mmu.Arl13b-GFP)* (Borovina et al., 2010)) and Cmcl2::GFP *(Tg(myl7:eGFP)* (Huang et al., 2003)) were used. Experiments were approved by the University of Bristol Ethical Review Committee and the UK Home Office.

### Morpholino knockdown

A translational block morpholino (Golgb1-ATG; AACATGGCTGACCTGCAAGAAAATA) and a splice site blocker morpholino at the boundary between the preceding intron and exon 14 (Golgb1-E14; CTGTTCCAGCTACTTATTGAAAAA) were obtained (Ensembl release #69) along with standard control morpholinos (Genetools LLC, Philomath, OR). 3.8 ng morpholino with 0.1M KCl and 0.1% phenol red was injected in one to four cell stage embryos. N-capped Rn-flagGolgb1 mRNA was *in vitro* transcribed (Ambion™ T7 mMessage mMachine kit, Fischer Scientific) from a pSG5 vector (SalI linearized, NEB) containing the template (gift from M.P. Lowe, University of Manchester, UK), and co-injected (375 pg) with MOs. The following primer sets were used for touchdown PCR (G-Storm) on cDNA (Superscript III, Invitrogen) from total RNA (RNeasy mini kit, cat# 74104, Qiagen), exon 13 forward (CCCAAAAGGAGAAGTGTGGA), exon 14 forward (AGATGCAAGTGCAACGGTCT), exon 15 reverse (ATTTTGATGCCTGTGCTTCC), and exon 16 reverse (GGGCAGCATCTAATGCAAGT). Morphants were scored in four classes (WT-like, mild, moderate, and severe) based on none, 1-2 mild, 3 with one severe, or 4 with at least two severe morphological characteristics respectively.

### *Golgb1* mutant zebrafish

F2 fish carrying the *sa11389* allele were acquired from the European stock centre (EZRC, Karlsruhe, Germany) and subsequently outcrossed with London AB WT fish (Kettleborough et al., 2013). F3 and F4 heterozygote in-crosses were used. For TALEN site directed mutagenesis (Bedell et al., 2012) the following optimised repeat variable domain (RVD) arm sequence (Cermak et al., 2011; Streubel et al., 2012) was cloned into a pGoldy vector from the Golden Gate TALEN and TAL effector Kit 2.0 (Kit #1000000024, Addgene): arm1 (upstream) NH HD HD NI NI HD NG HD NG NH HD HD NI HD HD HD NG HD NG and TALEN arm2 (downstream) NH NH NG NH NG NG NG NG NH HD NH NI NI HD NG NH NI NI NH. These flank the CCTCAGCTGTTTCAG spacer sequence containing a PVUII (NEB) restriction digest site. pGoldy-Golgb1-exon14-Arm1 and pGoldy-Golgb1-exon14-Arm2 constructs (SacI linearized) were *in vitro* transcribed to produce stable N-terminus capped mRNA (Ambion™ T3 mMessage mMachine kit, Fischer Scientific) and both arm transcripts (170 pg each) were injected in 1-cell stage embryo cytoplasm (London AB). F1 was screened for germline transmission by fin clips. Both mutant lines were genotyped by touchdown PCR using forward (5'-AGACAGGGTGCTTAGCCAAT-3') and reverse (5'-TGACAGCCTGATCTCTTGCA-3') primers. *Goigb1^X3078^* PCR product was PVUII digested to determine loss of restriction site whereas *golgb1^Q2948X^* PCR product was sequenced (MWG-Eurofins) using 5'-TCAATGCGGAGAATGCCAAG.

### *In vitro* tissue culture, short-interference RNA knockdown

Pig kidney epithelial cells (LLC-PK1) cells were maintained under standard conditions (37^o^C in 5% CO2) in DMEM (Medium-199, Sigma-Aldrich) plus 10% fetal calf serum (FCS). Calcium phosphate knockdown (Watson and Stephens, 2006) was performed with siRNA duplexes Si#C SsGolgb1 (GUUCAGUGAUGCUAUUCAA) and Si#D SsGolgb1 (UCACAUGUGUACCGAGGUA) (MWG-Eurofins, Germany).

Human telomerase immortalized retinal pigment epithelial cells (hTERT-RPE1, Takara Bioscience) were grown in DMEM F12 HAM supplemented with 10% FCS (Life Technologies, Paisley, UK). For giantin siRNA knockdown experiments, cells were first reverse transfected and then forward transfected 48 hours later using Lipofectamine 2000 according to the manufacturer's guidelines (Invitrogen, Carlsbad, CA). In each round of transfection 200 pmol each of siRNA duplex 1 (ACUUCAUGCGAAGGCCAAATT) and siRNA duplex 2 (AGAGAGGCUUAUGAAUCAATT) targeting giantin were pooled together or for mock transfections 400 pmol siRNA targeting GL2 (GUACGCGGAAUACUUCGAUU) was used.

### Immunofluorescence and histology

To achieve cilia formation, LLC-PK1 and hTERT-RPE1 cells were grown to confluence and serum starved for 24 hours prior to methanol fixation. Wholemount immunofluorescence and immunohistochemistry embryos or larvae were blocked in 10% FCS plus 1% bovine serum albumin blocking buffer. Primary antibodies: mouse α-acetylated tubulin (T6793, Sigma-Aldrich, UK), chicken α-GFP (GFP-1020, Aves Labs), rabbit polyclonal α-Collagen2a1 (ab34712, Abcam), mouse monoclonal α-chondroitin sulphate type A and C (CS-56, Sigma-Aldrich), mouse M2 α-flag (F1804, Sigma-Aldrich), sheep α-Golgin-84 (raised against coiled-coiled domains, gift from Martin Lowe, University of Manchester, UK). After primary antibody incubation samples were blocked again to reduce background staining. Secondary antibodies were highly cross-adsorbed Alexa-dye labelled anti-IgG (Thermo Fisher Scientific). Images were acquired using confocal microscopy using either a Leica SP5-II or SP8 AOBS confocal laser scanning microscope attached to a Leica DM I6000 inverted epifluorescence microscope. For cilia imaging a 63x lens (numerical aperture 1.3 for SP5 and 1.4 for SP5-II) with 5x zoom and argon laser were used.

For cell experiments, cells were washed in PBS, fixed in 100% MeOH at -20^o^C for 3 minutes, rinsed in PBS and blocked in 3% BSA-PBS for 30 minutes at room temperature. Antibodies were diluted in block and primary and secondary antibody labelling processed for 1 hour each, washing 3x in between. Finally, cells were labelled with DAPI ([4,6-diamidino-2-phenylindole (Life Technologies, Paisley, UK, D1306)]) and mounted in Mowiol (Calbiochem). Fixed cells were imaged using an Olympus IX70 microscope with 60x 1.42 NA oil-immersion lens, Exfo 120 metal halide illumination with excitation, dichroic and emission filters (Semrock), and a Photometrics Coolsnap HQ2 CCD, controlled by Volocity 5.4.1 (Perkin Elmer). Chromatic shifts in images were registration corrected using TetraSpek fluorescent beads (Thermo Fisher). Images were acquired as 0.2 μm z-stacks unless otherwise stated in the figure legend.

### Scanning electron microscopy

Both LLC-PK1 and pre-fixed (2 hours of 4% paraformaldehyde) 3 dpf larvae were fixed with 2.5% glutaraldehyde (100mM sodium-cacodylate) for 1 hour with subsequent 100mM sodium-cacodylate washes. Further fixation by 2% OsO_4_ (100mM sodium-cacodylate) for 3 hours. Dehydration steps: 25%, 50%, 70%, 80%, 90%, 96%, absolute, absolute over a course of 2 days. Critical point drying (Leica, EM CPD 300) was performed to remove excess of ethanol (12 cycles, 50% CO_2_ infusion speed). Sputter coating with gold palladium supplied with Argon gas (Emitech K575X) was performed prior to imaging under high vacuum.

### Reverse transcriptase PCR and quantitative Real-Time PCR

Larvae (5 dpf, n=30) were pooled and total RNA was isolated using RNeasy mini kit (cat# 74104, Qiagen) and reverse transcriptase reaction was performed by using Superscript III (cat# 18080093, ThermoFisher Scientific) according to the manufacturers' protocol. Quantitative Real-Time PCR (qPCR) reaction (primers: Table S1) was undertaken with DyNAmo HS SYBR green (F410L, ThermoFisher Scientific) cycling (40 times) at 95°C 25 seconds, 57.5°C 30 seconds, and 70°C 45 seconds followed by a standard melt curve (QuantStudio3, Applied Biosystems).

### Smoothened agonist, SHH and smoothened antagonist-1 treatment

Co-injected embryos with 3.8ng of control, ATG, or E14 MO with either 1ng control or p53^ATG^ MO (Chen et al., 2005) were treated with final concentrations of 0.15%v/v DMSO, 20 μM smoothened agonist (Sigma Aldrich), or 20 μM smoothened antagonist-1 from 4 hpf until 48 hpf. Embryos were dechorionated and put in fresh treatment solution at 24 hpf. Images were taken with a fluorescence stereomicroscope (Leica Microsystems, Mannheim, Germany).

For cell experiments, hTERT-RPE1 cells were subjected to two rounds of transfection. Six hours after the second round, cells were serum starved in un-supplemented DMEM F12 HAM overnight, then treated with either 500ng/ml recombinant SHH (C42II, N-terminus, R&D systems, Abingdon, UK, 1845-SH-025), 500ng/ml SANT1 (Sigma, Dorset, UK, S4572) or an equivalent volume of drug vehicle control for 24 hours before methanol fixation and staining.

## Acknowledgements

We also thank Martin Lowe for reagents and helpful discussions, Brian Ciruna and Tanya Whitfield for the supply of *Tg(Act-B2::Mmu.Arl13b-GFP)* transgenic fish, and members of the Hammond and Stephens' labs for continued input into this work. We would also like to thank Josi Peterson-Maduro, Ive Logister and Stefan Schulte-Merker for assistance with TALEN mutagenesis protocol. We thank the MRC and Wolfson Foundation for support of the Wolfson Bioimaging Facility and Judith Mantel for technical support.

## Author contributions

DJB and NLS designed and performed experiments, and analysed data; RES helped with the zebrafish experiments; DJS and CLH conceived and managed the project and contributed to data analysis. DJB wrote the paper with help from NLS, DJS and CLH.

---

**Figure S1:**
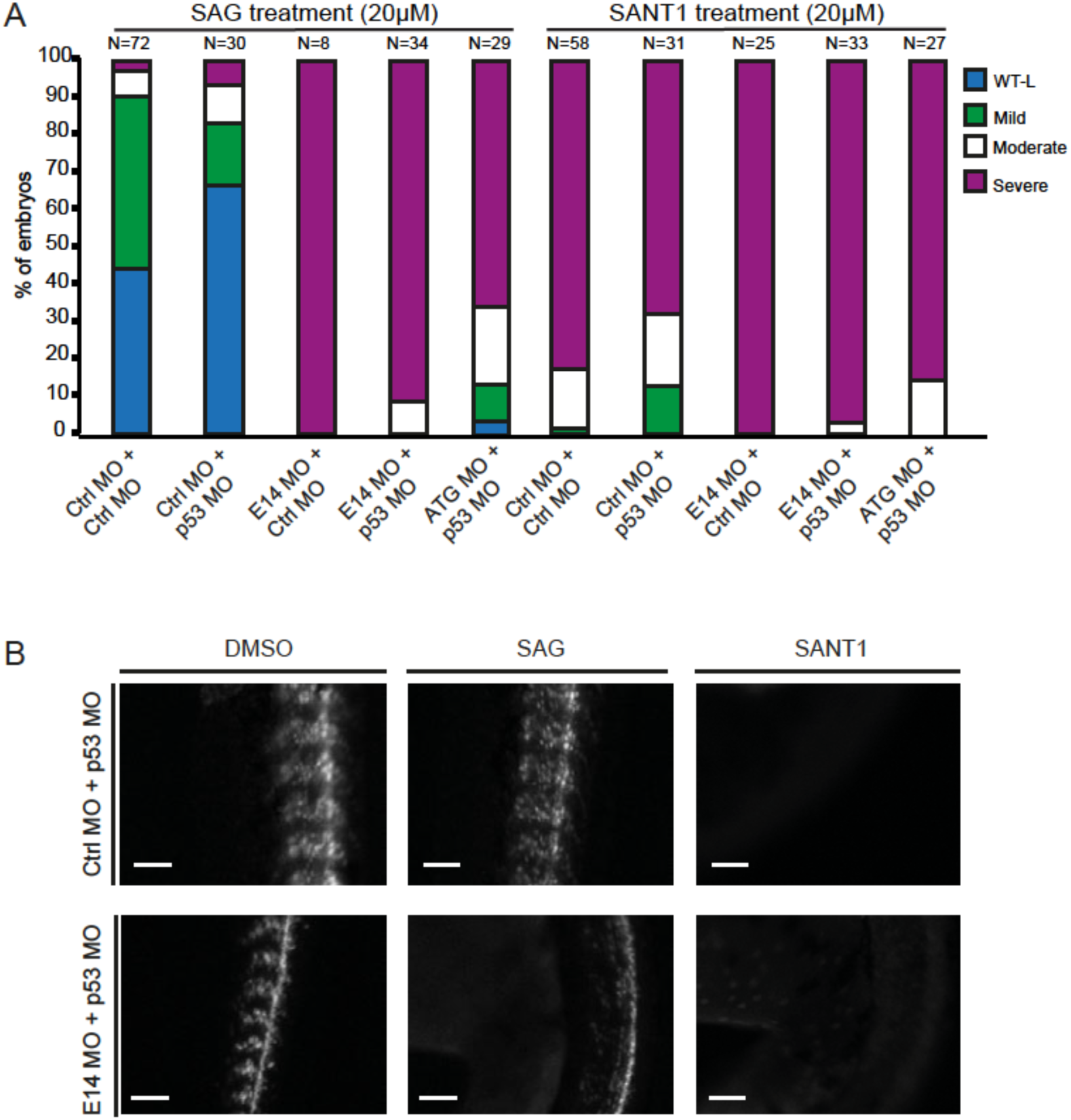
Smoothened agonist treatment neither alters the ciliopathy-like phenotypes in giantin morphants. (A) Quantification of phenotypes observed (48 hpf). Note, no difference is seen with 1 ng p53 morpholino co-injection in all conditions. (B) Gli::mCherry expression was reduced in somites after SANT1 treatment, but not in SAG-treated individuals (48 hpf). SAG: Smoothened agonist, SANT1: Smoothened antagonist 1.

**Figure S2:**
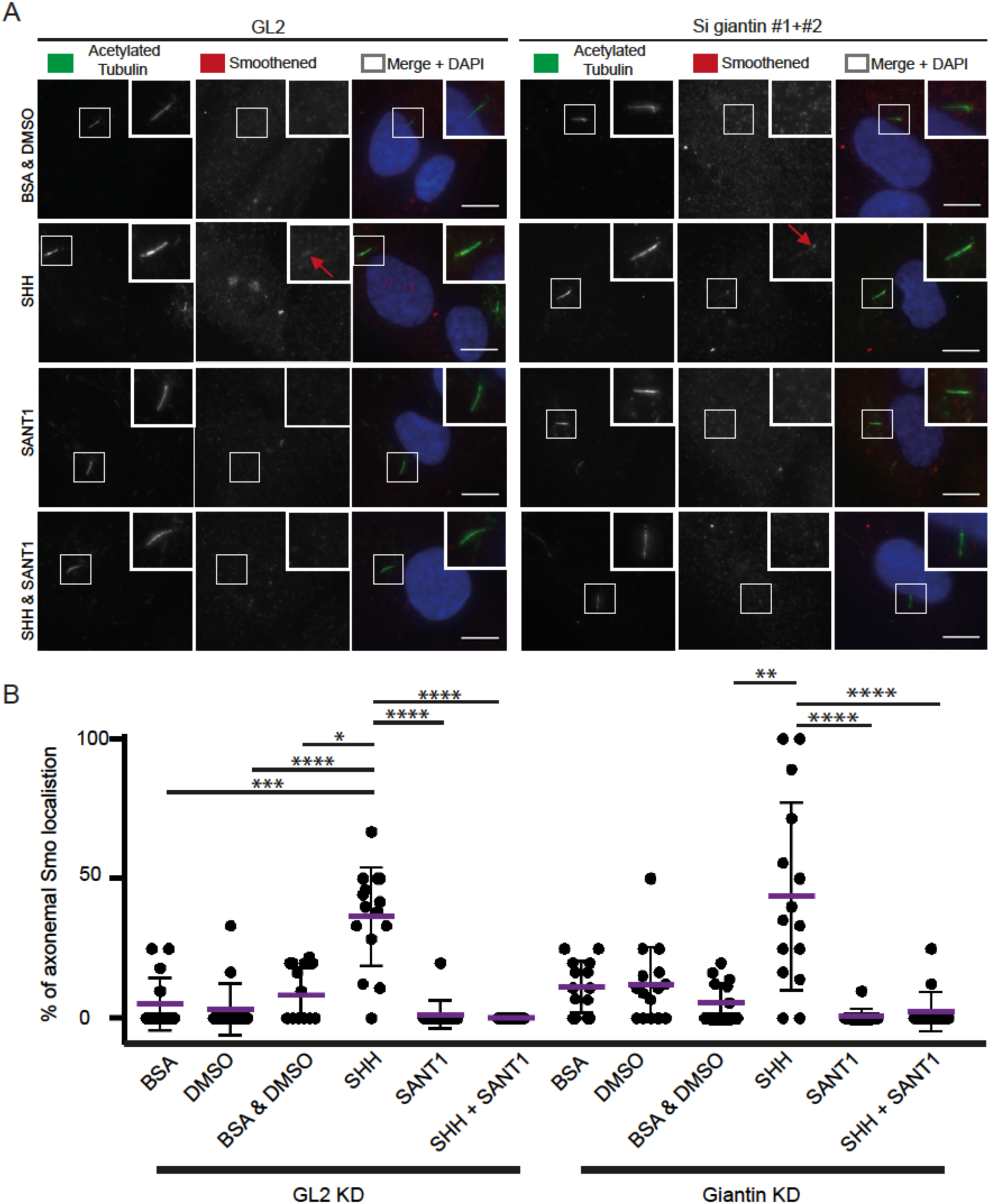
Depletion of giantin by RNAi in hTERT-RPE1 cells does not affect translocation of smoothened to the primary cilium. (A) Immunofluorescence of hTERT-RPE1 cells incubated with GL2 siRNA or two giantin siRNAs incubated for 96 hours, immunolabelled against acetylated Tubulin and Smo. Smoothened translocation was detected in both conditions upon Shh stimulation (red arrow). (B) Quantification of Smo translocation to the cilium of three independent experiments (5 fields per images per condition). Scale bar = 5 μm. SANT1: Smoothened antagonist 1, Shh: Sonic Hedgehog.

**Table S1:**
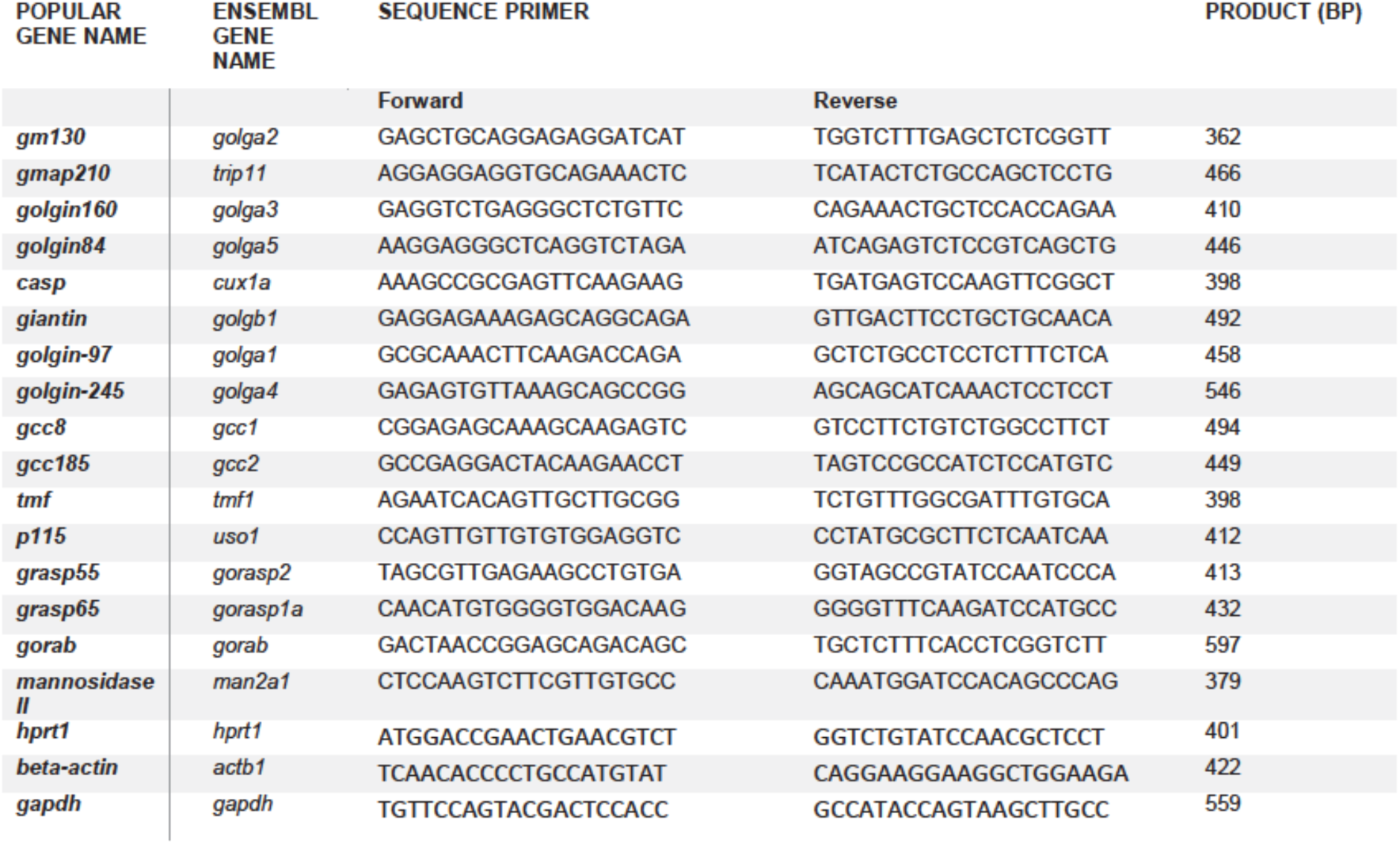
Primers used for quantitative real-time PCR of zebrafish golgins

